# Forward genetic screen for *Caenorhabditis elegans* mutants with a shortened locomotor healthspan

**DOI:** 10.1101/487876

**Authors:** Kazuto Kawamura, Ichiro N. Maruyama

## Abstract

Two people with the same lifespan do not necessarily have the same healthspan. One person may retain locomotor and cognitive functions until the end of life, while another person may lose them during adulthood. Unbiased searches for genes that are required to maintain locomotor capacities during adulthood may uncover key regulators of locomotor healthspan. Here, we take advantage of the relatively short lifespan of the nematode *Caenorhabditis elegans* and develop a novel screening procedure to collect mutants with locomotor deficits that become apparent in adulthood. After ethyl methanesulfonate mutagenesis, we isolated five *C. elegans* mutant strains that progressively lose adult locomotor activity. In one of the mutant strains, a nonsense mutation in Elongator Complex Protein Component 2 (*elpc-2)* causes a progressive decline in locomotor function. Mutants and mutations identified in the present screen may provide insights into mechanisms of age-related locomotor impairment and the maintenance of locomotor healthspan.

## Introduction

Locomotor ability indicates an animal’s healthspan across many species such as worms, flies, mice, and humans (Cesari et al., 2009; Grotewiel et al., 2005; Hahm et al., 2015; Justice et al., 2014). In these species, declines in locomotor capacities can be a feature of the normal aging process, or a symptom of an age-related disease. Currently, the genetic regulators that work to prevent age-related declines in locomotor function is largely unknown.

Recent studies have suggested that the genetic bases of lifespan and healthspan may not completely overlap (Bansal et al., 2015; Iwasa et al., 2010; Tissenbaum, 2012). From a candidate-based genetic screen, Iwasa *et al*. found that activation of the epidermal growth factor signaling pathway prolongs adult swimming ability in *C. elegans* without large effects on lifespan (Iwasa et al., 2010). More examples of genetic pathways that work to maintain locomotor healthspan may be discovered by carrying out unbiased searches for mutant animals that show progressive declines in locomotor capacity.

A forward genetic screen using *C. elegans* has previously been employed to identify genes that affect locomotor function during development (Brenner, 1974). However, unbiased screens that focus on locomotor deficits occurring later in life have not been carried out, in part due to the difficulty in distinguishing whether symptoms observed during adulthood were already present during development.

In the present study, we established the “Edge Assay” to measure locomotor ability of hundreds of adult worms at once. Using the Edge Assay, we developed a screening procedure to remove mutant worms with strong developmental locomotor defects on the first day of adulthood, and then isolated mutant worms that progressively lose their locomotor function on the third or fifth days of adulthood. After ethyl methanesulfonate-mutagenesis, we isolated five mutant strains that progressively lose their ability to complete the Edge Assay. In one mutant strain, we found that a mutation in the *elpc-2* gene causes progressive loss of locomotor function. *elpc-2* works with other Elongator complex genes, *elpc-1* and *elpc-3*, to maintain adult locomotor function in *C. elegans*. Along with the Elongator complex mutants, isolated mutants from our screen can be used as tools to explore mechanisms that work to maintain adult locomotor function in *C. elegans*.

## Results

### The “Edge Assay” can test locomotor function of hundreds of worms

Our forward genetic screen isolates mutant worms that progressively lose locomotor function. We established the Edge Assay to measure locomotor activity of hundreds of worms at once. The Edge Assay is carried out on a 9-cm agar plate with *E. coli* bacterial feed spread only on the outer edge of the plate. Up to a few hundred adult worms are placed on the center of the plate where there is no food (Fig. 1A; Fig. S1). Motile worms reach the *E. coli* on the edge of the plate, while worms with defects in locomotion or chemotaxis remain in the center of the plate.

**Figure 1.**
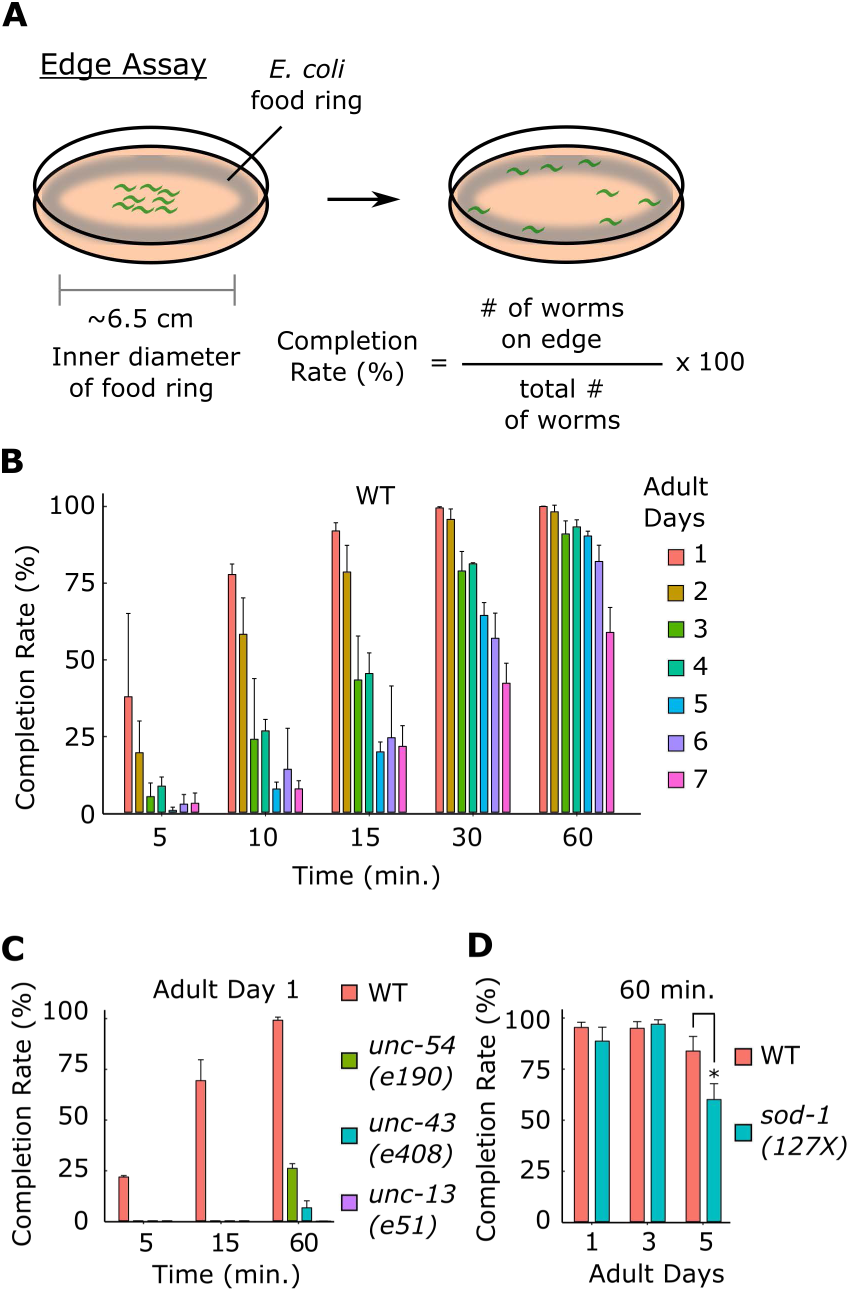
“Edge Assay” can measure locomotor ability of worms. (A) (*Left*) Schematic diagram of an Edge Assay plate immediately after placing worms at the center of the plate. (*Right*) Schematic diagram of Edge Assay plate after most worms reached the edge. (B) Edge Assay completion rates of wild-type worms from adult day 1 to 7 after 5, 10, 15, 30, and 60 min. (C) Completion rates for WT and developmental mutants deficient in locomotor function, *unc-54(e190)*, *unc-43(e408)*, and *(unc-13(e51)*. (D) Completion rates of WT and a previously reported *C. elegans* model of amyotrophic lateral sclerosis (Hsa-sod-1(127X)). For Edge Assay experiments, n = 3 biological replicate plates with each plate starting with approximately 100 worms per plate on adult day 1. Error bars indicate 95% confidence intervals. \**P* < 0.05; Unpaired Student’s *t* test for D.

On the first day of adulthood, 91.3% of wild-type worms reached the edge in 15 min and 99.6% reached the edge in 60 min (Fig. 1B; Fig. S1). *C. elegans* mutant strains that are defective in the function of neurons (*unc-13(e51)*, *unc-43(e408)*) (Maruyama and Brenner, 1991; Reiner et al., 1999) or muscles (*unc-54(e190)*) (MacLeod et al., 1981) could not reach the edge in 15 min on the first day of adulthood (Fig. 1C). After 60 min, 26% of *unc-54(e190)* mutants, 6.4% of *unc-43(e408)* mutants, and 0% of *unc-13(e51)* mutants reached the edge (Fig. 1C). Therefore, carrying out the Edge Assay for 15 min on the first day of adulthood can separate wild-type worms from worms with strong developmental locomotor defects.

On average, over 90% of wild-type worms could complete the Edge Assay in 60 min during the first five days of adulthood (Fig. 1B). A *C. elegans* model of amyotrophic lateral sclerosis (Hsa-sod-1(127X)) (Gidalevitz et al., 2009) showed a significant reduction in Edge Assay completion rate compared to wild-type worms on the fifth day of adulthood (Fig. 1D). Therefore, carrying out the Edge Assay for 60 min on the fifth day of adulthood can separate wild-type worms from worms that progressively lose their locomotor activity.

### Isolation of mutants that progressively lose locomotor activity during adulthood

We mutagenized wild-type N2 worms using ethyl methanesulfonate, and screened 3352 F2 offspring from 500 F1 worms (1000 genomes) (Table S1). We carried out the Edge Assay for the mutagenized F2 offspring on the first day of adulthood (Fig. 2A). To remove worms with developmental defects, worms that could not complete the Edge Assay in 15 min were aspirated away (Fig. 2A). Only worms that completed the Edge Assay on the first day of adulthood were kept for further screening. On the third and fifth days of adulthood, we tested the worms again with the Edge Assay and collected slow or uncoordinated mutants that remained near the center of the Edge Assay plate after 60 min (Fig. 2A). By removing worms with strong developmental defects on the first day of adulthood, we were able to isolate worms that progressively lost locomotor function during adulthood. We isolated 22 viable mutants, and created individual strains from those mutants (Table S1). Five of those mutant strains reproducibly showed progressive deficits in completing the Edge Assay during adulthood (Fig. 2B).

**Figure 2.**
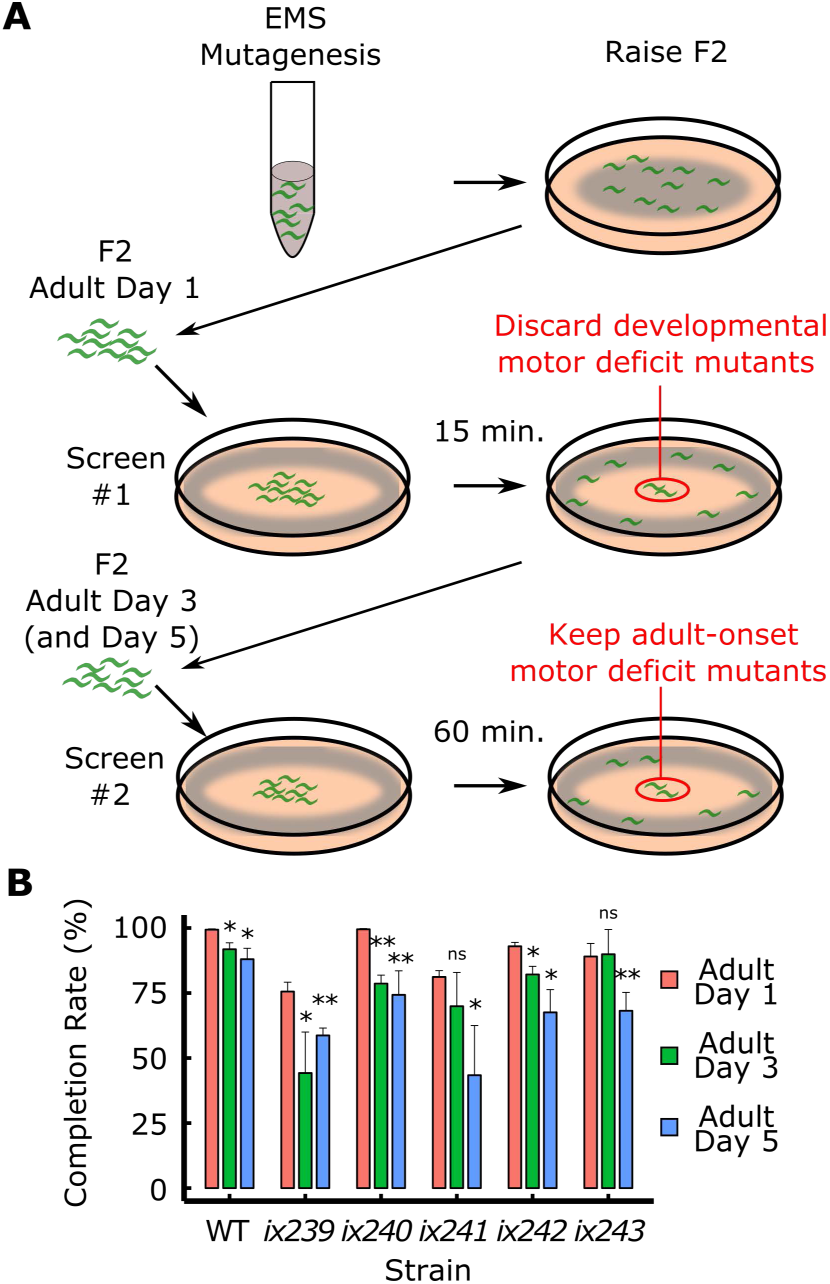
Isolation of mutant strains that progressively lose locomotor ability. (A) Schematic description of a forward genetic screen to isolate mutants that progressively lose locomotor ability. (B) Edge Assay completion rates of mutants identified from the screen. Error bars indicate 95% confidence intervals. n = 3 biological replicate plates, with each plate starting with approximately one hundred worms per plate on adult day 1. \**P* < 0.05; \*\**P* < 0.01; ns, not significant; Paired Student’s *t* test vs. adult day 1 completion rate.

To determine whether isolated mutant strains have deficits in locomotor function and not sensory function or search behavior, we measured locomotor function of worms on an agar plate without food. We recorded one-minute videos of 15 worms freely moving on a plate, and measured the maximum velocities and total travel distances for each worm. For each strain, we recorded three plates of 15 worms on the first, third, and fifth days of adulthood. All isolated mutant strains showed significantly greater reductions in maximum velocity and travel distance from the first to fifth days of adulthood compared to wild type except for *ix240* worms (Fig. 3A, B, D; Fig. S2A–D; Fig. S3A–D). In the *ix240* worms, progressive deficits other than locomotor function, such as sensory function or search behavior, may cause the reduction in Edge Assay completion rate.

**Figure 3.**
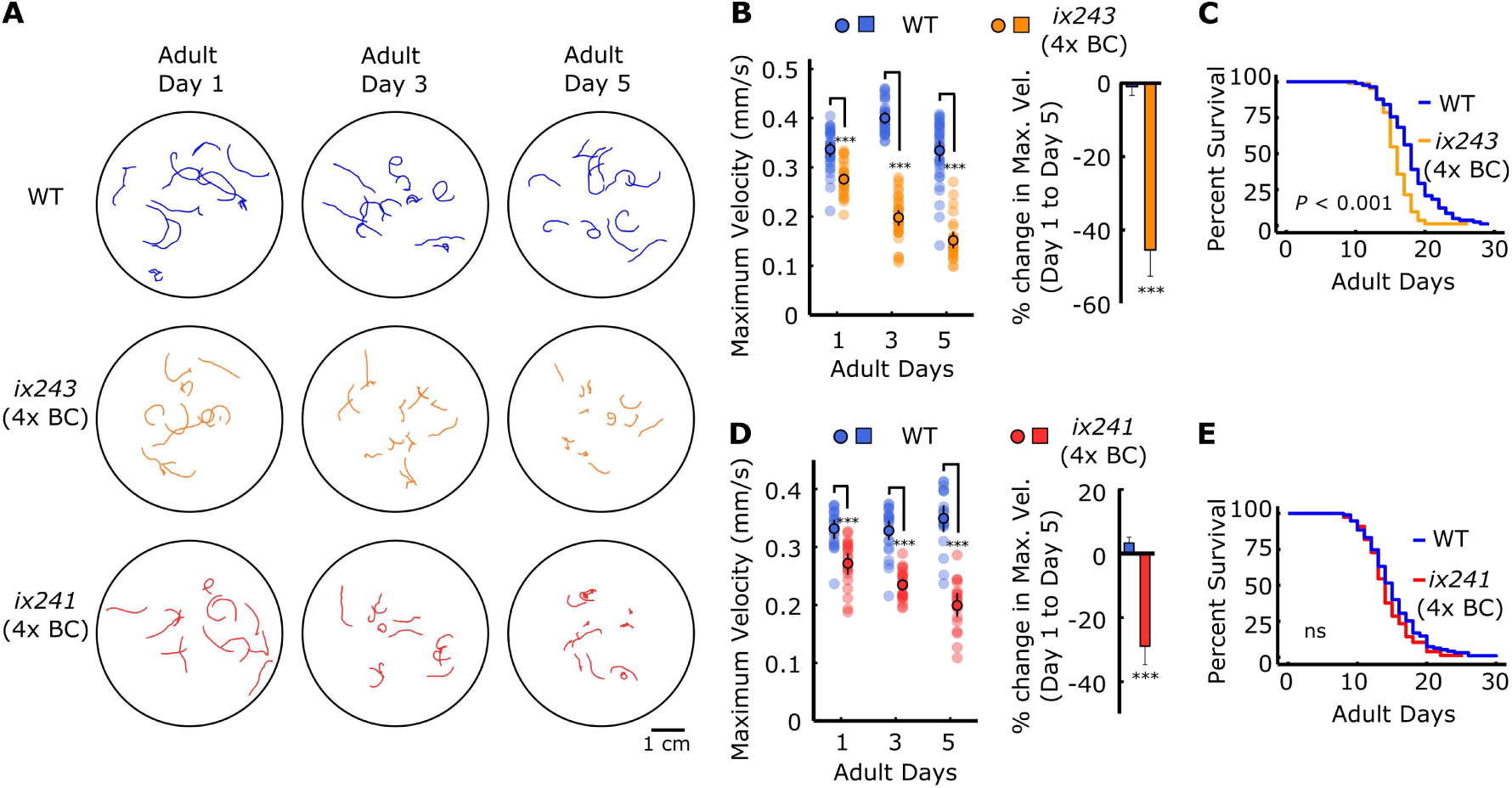
*ix241* and *ix243* worms show progressive locomotor decline after four backcrosses. (A) Representative locomotor tracks from 1-min video recordings of wild-type (WT), *ix243*(backcrossed four times (4x BC)), and *ix241*(4x BC) worms on adult days 1, 3, and 5 on plates with no food. n = 10–15 tracks per plate (some worms were unable to be tracked for a full minute, and were removed from analysis) (B) (*Left*) Maximum velocities of WT and *ix243*(4x BC) worms. (*Right*) Percent change in maximum velocity of WT and *ix243*(4x BC) worms on adult day 5 compared to adult day 1. (C) Survival curve of WT (n = 56 worms) and *ix243*(4x BC) (n = 89) worms.(D) (*Left*) Maximum velocities of WT and *ix241*(4x BC) worms. (*Right*) Change in maximum velocity of WT and *ix241*(4x BC) worms. (E) Survival curve of WT (n = 94) and *ix241*(4x BC) (n = 77) worms. Error bars indicate 95% confidence intervals. For maximum velocity experiments, n = 30–45 worms per strain for each day (10–15 worms from 3 biological replicate plates). For percent change in maximum veloity graphs, n = 3 biological replicate plates. \*\*\**P* < 0.001; ns, not significant; Unpaired Student’s *t* test for maximum velocity comparisons; Log-rank test for lifespan comparisons.

### *ix241* and *ix243* mutant strains show progressive decline in locomotor function

*ix241* and *ix243* worms were backcrossed with the parental N2 strain to reduce the number of mutation sites that do not affect locomotor function. After each backcross, we checked for individual lines that still showed a progressive decline in locomotor function. We measured the maximum velocity and travel distance of individual worms on an agar plate without food on the first, third, and fifth days of adulthood. *ix241* and *ix243* worms still showed significant reductions in both maximum velocity and travel distance after the fourth backcross (Fig. 3A, B, D; Fig. S3A–D).

To check whether *ix241* and *ix243* worms were simply aging faster than wild-type worms, we measured lifespans of the two strains. The lifespan of *ix241* worms was not significantly shortened compared to that of wild type (Fig. 3E; Table S2). The median lifespan of the *ix243* worms was shortened by two days (Fig. 3C; Table S2). To compare relative reductions in lifespan and locomotor healthspan, we measured the maximum velocities of wild type and *ix243* worms for 10 days (Fig. S4A). We quantified the percent decrease in lifespan by comparing the areas under the survival curves of wild type and *ix243* worms (Fig. S4B). We quantified the percent decrease in locomotor healthspan by comparing the areas under the decline in maximum velocity curves of wild type and *ix243* worms (Fig. S4C). For *ix243* worms, there is an average 11.5% reduction in lifespan, while there is a significantly greater 18.5% reduction in locomotor healthspan (Fig. S4D).

*ix243* worms take a 13.9% longer time to reach adulthood (Table S3). The developmental delay was taken into account for locomotor and lifespan measurements by allowing *ix243* worms an extra 10 h to develop, and starting locomotor and lifespan measurements from the first day of adulthood. *ix243* worms show a 17.7% decrease in maximum locomotor activity on the first day of adulthood compared to wild-type worms (Fig. 3B). The deficit in locomotor capacity compared to wild-type worms increases to 54.8% on the fifth day of adulthood (Fig. 3B). These results suggest that the *ix243* mutant allele has modest negative effects on development and lifespan, with relatively stronger negative effects on locomotor healthspan.

*ix241* worms take 4.0% longer to reach adulthood (Table S3) and show an 18.1% decrease in maximum locomotor activity on the first day of adulthood compared to wild-type worms (Fig. 3D). The deficit in locomotor function compared to wild-type worms increases to 43.0% on the fifth day of adulthood (Fig. 3D). The *ix241* mutant allele has no negative effect on lifespan, a modest negative effect on development, and a relatively stronger negative effect on locomotor healthspan.

### Nonsense mutation in *elpc-2* causes progressive loss of adult locomotor function in *ix243* worms

We used whole genome sequencing and a modified version of the sibling subtraction method to identify the causative mutation site in the *ix243* strain (Fig. S5) (Joseph et al., 2018). Mutations were evenly induced on all chromosomes in the *ix243* mutant strain before backcrossing (Fig. 4A). Many mutations remained on Chromosome III after comparing mutations in backcrossed strains that show a progressive loss of adult locomotor function and subtracting mutations in backcrossed strains that do not show progressive loss of adult locomotor function (Fig. 4B; Table S4). A nonsense mutation from TGG to TAG within the protein coding region of *elpc-2* was predicted to disrupt protein function (Fig. 4C; Table S4). Presence of the *elpc-2* mutation site was confirmed by Sanger sequencing (Fig. 4D).

**Figure 4.**
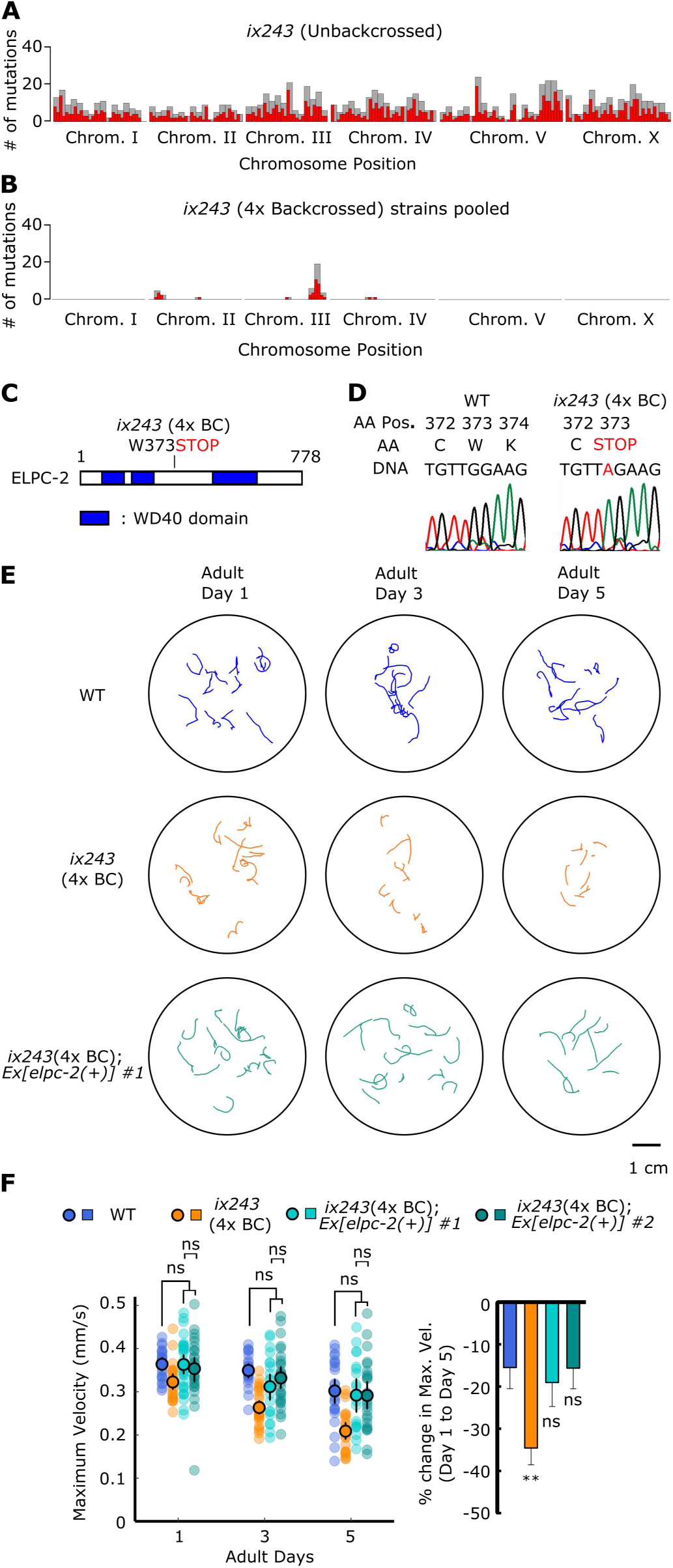
*elpc-2* mutation causes locomotor deficits in the *ix243* mutant strain. (A) Mutation frequency along each chromosome of *ix243* mutant strain before backcrossing. Red bars indicate 0.5-Mb bins and grey bars indicate 1.0-Mb bins. (B) Mutation frequency along each chromosome for pooled *ix243*(4x BC) worms. (C) Schematic diagram of ELPC-2 protein and location of mutation site in *ix243* allele. (D) *ix243* mutation site on ELPC-2 amino acid (AA) sequence and *elpc-2* DNA sequence. (E) Representative locomotor tracks of WT, *ix243*(4x BC), and *ix243*(4x BC);*Ex[elpc-2(+)]* #1 worms. (F) (Left) Maximum velocities of WT, ix243(4x BC), ix243(4x BC);Ex[elpc-2(+)] #1, and ix243(4x BC);Ex[elpc-2(+)] #2 worms. n = 30–45 worms per strain for each day (10–15 worms from 3 biological replicate plates). (Right) Percent change in maximum velocity of worms from left panel. n = 3 biological replicate plates. Error bars indicate 95% confidence intervals. **P < 0.01; ns, not significant; One-way ANOVA with Dunnett’s post hoc test vs. WT.

To test whether loss of *elpc-*2 causes a progressive decline in locomotor function, we injected a genomic fragment of *elpc-2* including 2090-base pairs (bp) upstream of the start codon and 851-bp downstream of the stop codon in the *ix243* mutant strain. The wild-type *elpc-2* fragment rescued the progressive loss of adult locomotor function (Fig. 4E, F; Fig. S6A, B). These results suggest that *elpc-2* is required for maintenance of adult locomotor function in *C. elegans*. The *ix243* mutant strain is the first reported mutant of the *elpc-2* gene in *C. elegans*.

### The Elongator complex is required to maintain locomotor function

ELPC-2 is a component of the Elongator complex. In *C. elegans*, there are four predicted components of the Elongator complex (ELPC1–4) (Solinger et al., 2010). To test whether functional loss of *elpc-2* causes the locomotor defect independently or as part of the Elongator complex, we measured locomotor activity of strains carrying deletions in *elpc-1* and *elpc-3*. We found that *elpc-1(tm2149)* and *elpc-3(ok2452)* mutant strains also cannot maintain locomotor function during adulthood (Fig. 5A, B). *elpc-1(tm2149);elpc-2(ix243)* and *elpc-2(ix243);elpc-3(ok2452)* double mutants did not show additive deficiencies in locomotor function (Fig. 5C; S7A–H). These results suggest that proper functioning of the entire Elongator complex is necessary to maintain locomotor healthspan. We assessed the expression pattern of *elpc-2* by creating an *elpc-2*p*::GFP* transcriptional reporter that expresses GFP under control of the *elpc-2* promoter. The transcriptional reporter was broadly expressed in many tissues including head and body wall muscles, head neurons, pharynx, canal cell, coelomocytes, intestine, and tail (Fig. S8A–C). The expression pattern of *elpc-2* overlaps with previously reported expression of *elpc-1* in the pharynx, head neurons, and body wall muscles (Chen et al., 2009).

**Figure 5.**
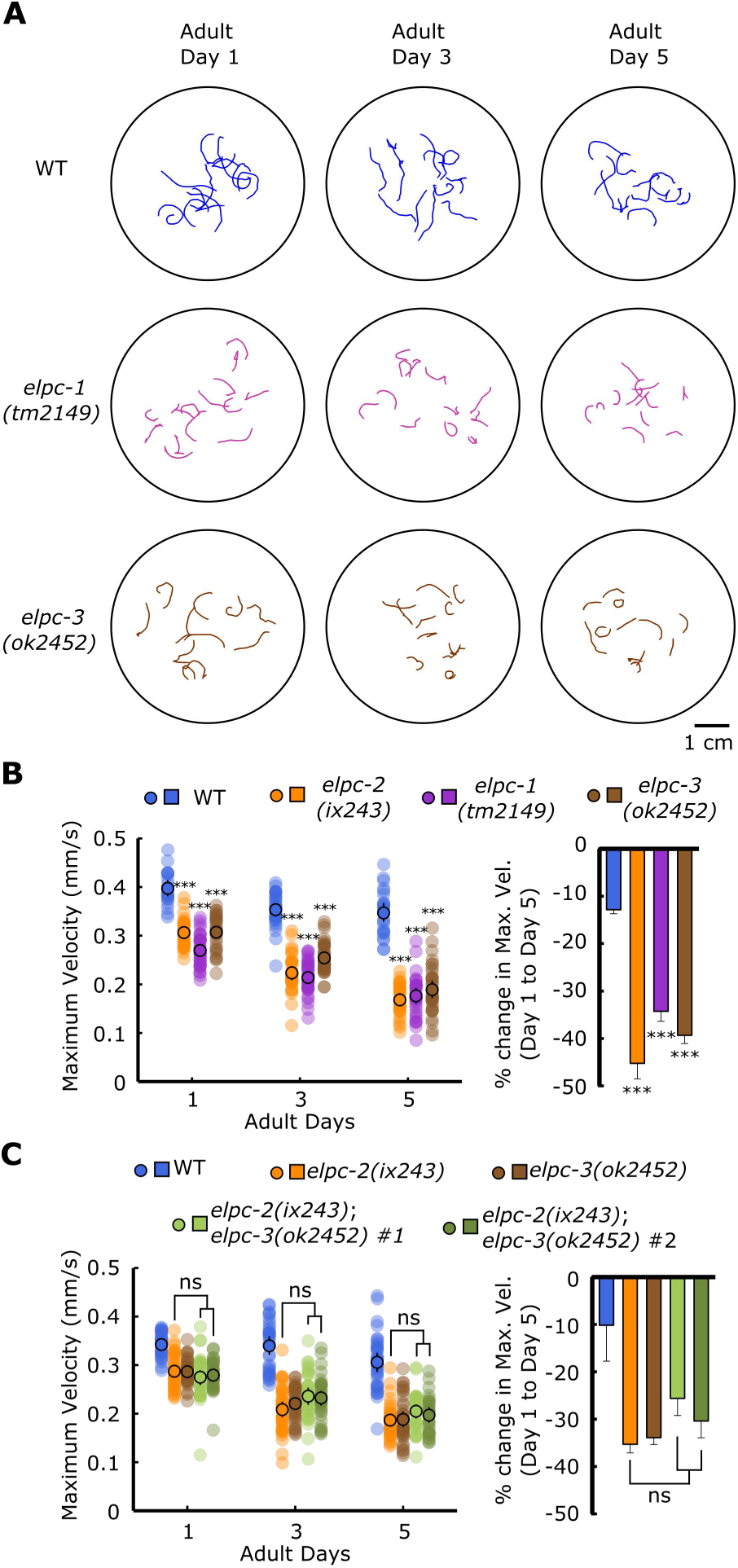
The Elongator complex is required to maintain locomotor function. (A) Representative locomotor tracks of WT, *elpc-1(tm2149)* and *elpc-3(ok2452)* worms. n = 10–15 tracks per plate. (B) (*Left*) Maximum velocities of WT *elpc-1(tm2149)* and *elpc-3(ok2452)* worms. (*Right*) Percent change in maximum velocity of worms from left panel. (C) (*Left*) Maximum velocities of WT, *elpc-2(ix243)*, *elpc-3(ok2452)*, and *elpc-2(ix243);elpc-3(ok2452)* worms. (*Right*) Percent change in maximum velocity of worms from left panel. Error bars indicate 95% confidence intervals. For maximum velocity experiments, n = 30–45 worms per strain for each day (10–15 worms from 3 biological replicate plates). For percent change in maximum veloity graphs, n = 3 biological replicate plates. \*\*\**P* < 0.001; ns, not significant; One-way ANOVA with Dunnett’s post hoc test vs. WT for B; One-way ANOVA with Tukey’s post hoc test for C.

### Loss-of-function mutation in *tut-1* also causes progressive decline in locomotor function

Mutants for *elpc-1* and *elpc-3* have previously been reported to modify the wobble uridine (U_34_) of tRNA by adding carbamoylmethyl (ncm) and methoxycarbonylmethyl (mcm) side chains to the 5’carbon of U_34_ (Chen et al., 2009; Nedialkova and Leidel, 2015). Wobble uridines with the mcm^5^ modification are further modified by TUT-1 to add a thio-group at the 2’ carbon to create mcm^5^s^2^U (Chen et al., 2009). In wild-type worms, only ncm^5^ and mcm^5^s^2^ modifications are present (Chen et al., 2009). In *tut-1(tm1297)* mutants, an mcm^5^ modification was observed, which is not normally present in wild-type worms (Chen et al., 2009). In *elpc* mutants, an s^2^ modification was observed, which is not normally present in wild-type worms (Chen et al., 2009).

To check whether loss of tRNA thiolation could cause a progressive decline in locomotor function, we measured the locomotor function of *tut-1(tm1297)* mutant worms. *tut-1(tm1297)* mutant worms showed a significantly greater decline in locomotor function during adulthood compared to wild-type worms, indicating that tRNA modifications may be a general mechanism involved in maintenance of locomotor healthspan in *C. elegans* (Fig. 6A; Fig. S9A, B).

**Figure 6.**
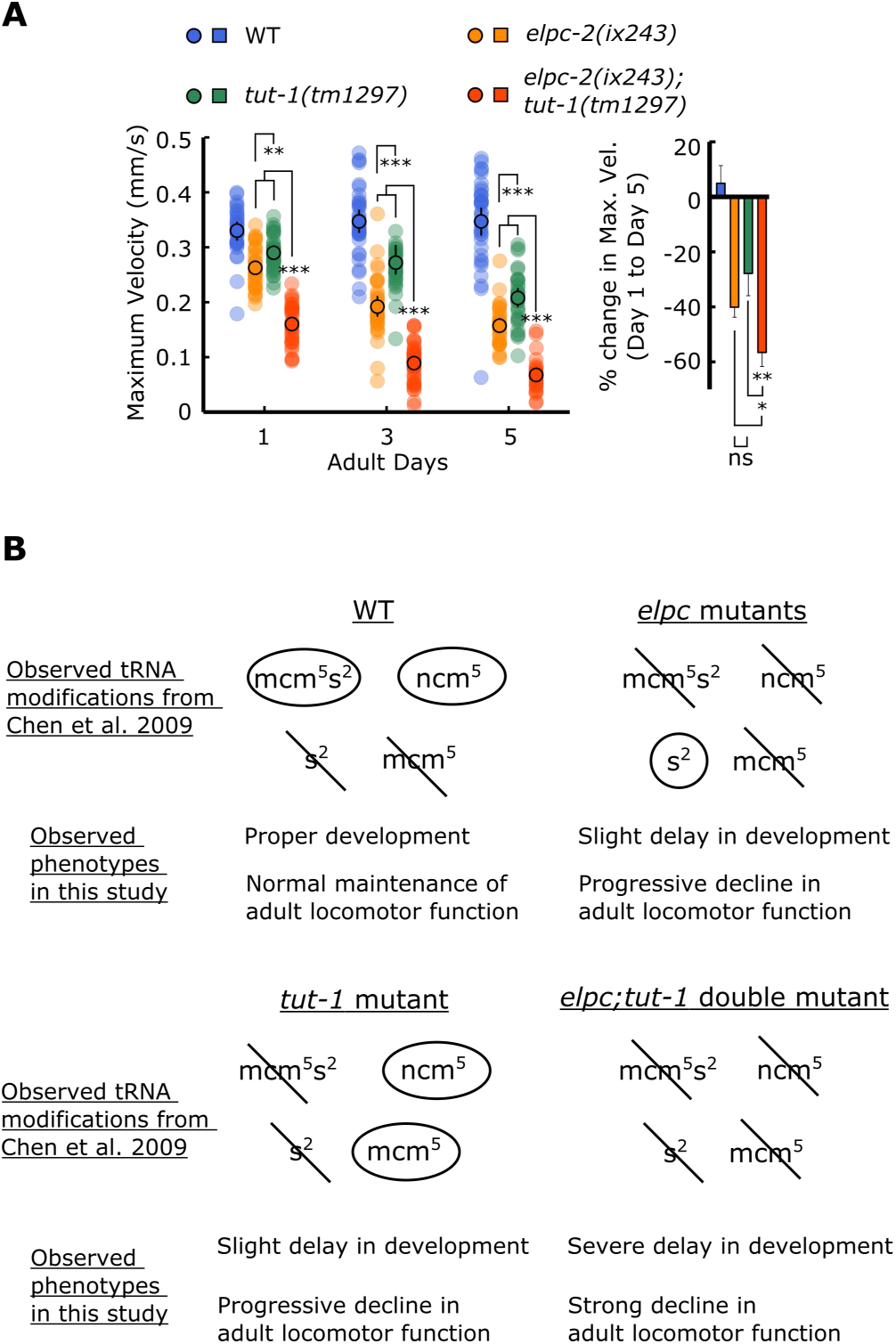
*tut-1(tm1297)* mutant shows progressive decline in locomotor function. (A) (*Left*) Maximum velocities of WT, *elpc-2(ix243)*, *tut-1(tm1297)*, and *elpc-2(ix243);tut-1(tm1297)* worms. (*Right*) Percent change in maximum velocity of worms from left panel. Error bars indicate 95% confidence intervals. n = 30–45 worms per strain for each day (10–15 worms from 3 biological replicate plates). For percent change in maximum velocity graphs, n = 3 biological replicate plates. \**P* < 0.05; \*\*\**P* < 0.001; ns, not significant; One-way ANOVA with Tukey’s post hoc test. (B) Summary of observed tRNA modifications in *elpc* and *tut-1* mutants from Chen et al. 2009, and summary of observed phenotypes in *elpc* and *tut-1* mutants from this study.

The *elpc-2(ix243)*;*tut-1(tm1297)* double mutant showed synthetic effects for locomotor function and for developmental maturation. *elpc-2(ix243)*;*tut-1(tm1297)* double mutant worms showed a strong defect in locomotor function on the first day of adulthood and a significantly greater reduction in maximum velocity and travel distance during adulthood relative to either of the single mutants (Fig. 6A; Fig. S9A, B). In addition, *elpc-2(ix243)*;*tut-1(tm1297)* double mutant worms took almost twice as long to reach adulthood (145.4 h) compared to *elpc-2(ix243)* worms (80.2 h) or *tut-1(tm1297)* worms (82.0 h) (Table S3; Table S5). The synthetic effects may be explained by the complete absence of U_34_ modifications in the *elpc-2(ix243)*;*tut-1(tm1297)* double mutant strain. The presence of the s^2^ modification in the *elpc* mutants, and the presence of the mcm^5^ and ncm^5^ modifications in the *tut-1* mutant may enable partial tRNA functionality and allow relatively proper development and partial capacities to maintain locomotor function (Fig. 6B).

## Discussion

In this study, we established the Edge Assay to simultaneously measure locomotor function of up to a few hundred adult worms. For our forward genetic screen, we used the Edge Assay to remove worms with strong developmental locomotor defects, and isolated worms with locomotor deficits that become apparent in adulthood. By carrying out the Edge Assay on the first day of adulthood, we were able to remove worms with strong developmental locomotor defects and overcome the difficulty of distinguishing developmental and progressive locomotor deficit mutants.

The Edge Assay may be used for a variety of applications involving locomotor function. For example, the Edge Assay can be used in suppressor screens to search for mutant worms that show improvements in locomotor function of previously characterized *C. elegans* models of neurodegenerative disease. It may also be possible to use the Edge Assay to screen for other types of progressive declines in functional capacity such as sensory or cognitive deficits by replacing the food ring with specific chemicals or learned cues.

The *ix241* and *ix243* mutant strains show similar declines in locomotor function, but have different phenotypes in regard to lifespan. This suggests that genes that regulate lifespan and locomotor healthspan may not completely overlap. In terms of improving quality of life, genetic regulators of healthspan may be better therapeutic targets than regulators of lifespan. Further studies and genetic screens that focus on healthspan-related phenotypes may provide novel insights into mechanisms that regulate healthspan and quality of life across many species.

In the *ix243* mutant strain, we found that *elpc-2* is required to maintain locomotor healthspan, and works as part of the Elongator complex. The Elongator complex is an evolutionarily conserved protein complex that consists of six subunits in *S. cerevisiae*, *A. thaliana*, *M. musculus*, and humans (Creppe and Buschbeck, 2011; Dauden et al., 2017). ELP1–ELP3 form the core complex, and ELP4–ELP6 form a sub-complex (Creppe and Buschbeck, 2011). In *C. elegans*, there are currently only four predicted homologs of the Elongator complex (*elpc-1*–*4*). From the present study, loss-of-function mutations in *elpc-1*, *elpc-2*, and *elpc-3* caused a shortened locomotor healthspan. Proper functioning of the Elongator complex may require multiple or all components of the complex (Dauden et al., 2017). The present work suggests that the Elongator complex is essential in maintenance of locomotor healthspan.

Allelic variants of ELP3, the catalytic subunit of the Elongator complex, were found to be associated with amyotrophic lateral sclerosis (ALS) in three human populations (Simpson et al., 2009). Risk-associated alleles have lower levels of ELP3 in the cerebellum and motor cortex of ALS patients, and protection-associated alleles have higher levels of ELP3 (Simpson et al., 2009). Overexpression of ELP3 reduced levels of axonopathy in the SOD1^A4V^ zebrafish model of ALS and SOD1^G93A^ mouse model of ALS (Bento-Abreu et al., 2018). The present work complements studies that have been performed in the context of ALS, and suggest that loss of the Elongator complex alone can cause locomotor deficits during adulthood in *C. elegans*. Future therapies that target multiple subunits of the Elongator complex may provide more robust effects than therapies that target only the catalytic ELP3 subunit.

ELP2 mutations were reported as the causative mutations in a familial form of neurodevelopmental disability (Cohen et al., 2015). Patients who are compound heterozygotes for two different ELP2 missense mutations demonstrate a lack of motor control starting in early development, severe intellectual disability, and progressive loss of locomotor function (Cohen et al., 2015). In our newly isolated *elpc-2(ix243)* strain, we also see deficits in locomotor function on the first day of adulthood and a delay in development (Fig. 3A, B; Fig. S3A, B; Table S3). Since the amino acid sequences of *C. elegans* ELPC-2 and human ELP2 are highly conserved (Fig. S10), some aspects of the neurodevelopmental dysfunctions that result from the human ELP2 mutation may be modeled in our *elpc-2(ix243)* mutant strain.

The Elongator complex was originally identified as a transcriptional regulator associated with RNA polymerase II (Otero et al., 1999). However, follow-up studies have found that the main functions of the Elongator complex may involve tRNA modification (Chen et al., 2009; Huang et al., 2005), and tubulin acetylation (Solinger et al., 2010). The tRNA thiolation mutant, *tut-1(tm1297)*, also showed a progressive decline in locomotor function. In yeast, tRNA modifications are important for proper translation and folding of proteins (Nedialkova and Leidel, 2015). tRNA modifications may affect locomotor healthspan by regulating translation efficiency and protein folding.

Starting from an unbiased forward genetic screen using *C. elegans*, we have found that mutations in Elongator complex and *tut-1* cause progressive declines in locomotor function during adulthood. Future screening procedures that utilize the Edge Assay, and further analysis of the isolated mutants from the present screen may provide insights into how locomotor function is maintained during adulthood.

## Methods

### Strains

*C. elegans* Bristol N2 strain was used as wild type. Worms were cultivated on Nematode Growth Media (NGM) agar plates with *E. coli* strain OP50 at 20°C (Brenner, 1974). See Supplementary Information Appendix for strains used in the present study.

### Edge Assay

Edge Assay plates were prepared by pouring 16 mL of NGM agar into a circular 9 cm plate. NGM plates were dried overnight with the lid on at 25°C, then kept at 4°C until use. On the day before the Edge Assay, a total of 100 µL of *E. coli* suspension was spotted on four spots near the edge of the NGM plate. The tip of a 50 mL serological pipette was briefly placed over a flame to smoothen the tip. The NGM plate was placed on an inoculating turntable and the smoothened pipette tip was held against the *E.coli* drop. The plate was slowly rotated while holding the pipette tip still. The plate was rotated 360° to spread the *E. coli* around the edge of the whole plate. Plates were incubated overnight at 25°C and used the next day. Synchronized worms were collected and washed twice with M9 buffer containing 0.1% gelatin. Worms were placed on the center of an Edge Assay plate and excess M9 buffer was removed with the edge of a Kimwipe. The number of worms that reached or did not reach the edge were counted at various time points to measure the Edge Assay completion rate.

### Isolation of mutants that show a progressive decline in locomotor function

Wild-type N2 worms were mutagenized and cultured as previously described (Brenner, 1974). Larval stage 4 worms were mutagenized by incubation in a 50 mM ethyl methanesulfonate solution for 4 h. EMS-mutagenized F2 adult day 1 worms were collected and washed twice with M9 buffer containing 0.1% aqueous gelatin. Worms were placed at the center of an Edge Assay plate and excess buffer was removed with the edge of a Kimwipe. After 15 min, worms that did not reach the edge were removed using an aspirator. Worms that reached the edge were maintained on the same plate until adult day 3. On adult day 3, worms were collected and washed with M9 buffer containing 0.1% gelatin and the Edge Assay was repeated on a new Edge Assay plate. Worms that were unable to reach the edge were collected as adult day 3 progressive locomotor deficit mutants. Worms that reached the edge were maintained on the same plate until adult day 5. On adult day 5, worms were collected and washed with M9 buffer containing 0.1% gelatin and the Edge Assay was repeated on a new Edge Assay plate. Worms that were unable to reach the edge were collected as adult day 5 progressive locomotor deficit mutants.

### Measurements of maximum speed and travel distance

Worms were synchronized by placing five adult day 1 worms onto an NGM plate with food, and allowed to lay eggs for 3 h. When the offspring reached adult day 1, 15 worms were picked randomly onto a 6 cm NGM plate without bacteria. After the worms moved away from the initial location with residual food, worms were again moved onto a different NGM plate without bacteria. Movement of worms was recorded for 1.0 min with a charge-coupled device camera INFINITY3-6URM (Lumenera Corporation, Ottawa, Canada). Images were analyzed using ImageJ and wrMTrck software (www.phage.dk/plugins) to produce maximum speed and travel distance (Nussbaum-Krammer et al., 2015). Measurements were made with the lid on in a temperature-controlled room set at 20 °C. At least three biological replicate plates of 15 worms each were measured for each strain. Worms that were lost during the video recording were not included in the analysis.

### Whole-genome DNA sequencing

*C. elegans* DNA was sequenced using the MiSeq platform (Illumina, San Diego, CA). Libraries were prepared with an Illumina TruSeq Library Prep Kit. Mapping was conducted with BWA software (Li and Durbin, 2009). Resulting files were converted to bam files, then to pileup format with Samtools (Li et al., 2009). Variant analysis was conducted using VarScan and SnpEff available on the Galaxy platform (Blankenberg et al., 2010; Cingolani et al., 2012; Giardine et al., 2005; Goecks et al., 2010; Koboldt et al., 2009). Mutation frequencies along the chromosome were calculated and visualized using CloudMap (Minevich et al., 2012).

### Transcriptional reporter expression

A genomic fragment of 2090-bp immediately upstream of the start codon of the *elpc-2* gene was PCR-amplified using “5’ *elpc-2*p overlap ppd95.79 107-” and “3’ *elpc-2*p overlap ppd95.79 138-” primers, which have 15-bp overhangs that anneal immediately upstream of the GFP sequence in the pPD95.79 vector (See Supplementary Information for primers used). The pPD95.79 vector containing GFP was linearized by PCR using the “5’ ppd95.79 107-” and “3’ ppd95.79 138-” primers. The template vector was digested with restriction enzyme *Dpn*I (New England Biolabs, Ipswich, MA), and the linearized vector was purified by Wizard SV Gel and PCR Clean-Up System (Promega, Madison, WI). The pure linearized vector and the *elpc-2* promoter were fused using an In-Fusion HD Cloning Kit (Takara, Kusatsu, Japan) to make the *elpc-2*p*::GFP* transcriptional reporter construct. The construct was microinjected into the gonads of wild-type worms at a concentration of 50 ng/µL. Worms that expressed the reporter construct were immobilized in 25 mM sodium azide and observed under a confocal microscope LSM710 (Carl Zeiss, Oberkochen, Germany). A z-stack image was created from images taken at 1 µm increments.

### Creation of double mutants

Double mutant strains were created by crossing males of one strain with hermaphrodites of another. Double mutants were checked by extracting their DNA, amplifying a genomic fragment flanking the mutation site by PCR, and sequencing the PCR product by Sanger sequencing. See Supplementary Information for primer details.

### Statistics

All results are expressed as means with a 95% confidence interval. Student’s *t* test was used for pairwise comparisons with Excel 2010 (Microsoft). For multiple comparisons, one-way ANOVA was followed with Dunnett’s post hoc test or Tukey’s Honest Significant Difference test using R (Team, 2015). Statistical significance was set at \**P* < 0.05; \*\**P* <0.01; \*\*\**P* < 0.001.

## Data availability

All isolated strains and plasmids are available upon request. Whole genome DNA sequencing data is available on NCBI Sequence Read Archive (PRJNA530333 SAMN11311296 SAMN11311297).

## Acknowledgements

We thank H. Goto, M. Kanda, M. Kawamitsu, S. Yamasaki and other DNA sequencing section members for technical assistance with whole genome sequencing. We are grateful to T. Murayama and E. Saita for helpful advice and support. We thank D. Van Vactor, B. Kuhn and members of the Maruyama unit for helpful discussions and comments. We are grateful to H. Ohtaki for administrative support. We thank Steve Aird for technical editing of this manuscript. We thank the *Caenorhabditis* Genetics Center, which is funded by NIH Office of Research Infrastructure Programs (P40 OD010440), for providing worm strains. We also thank the National Bioresource Project (Japan) for providing worm strains. K. K. was supported by Japan Society for the Promotion of Science KAKENHI (Grant 16J06404).

---

**Figure S1.**
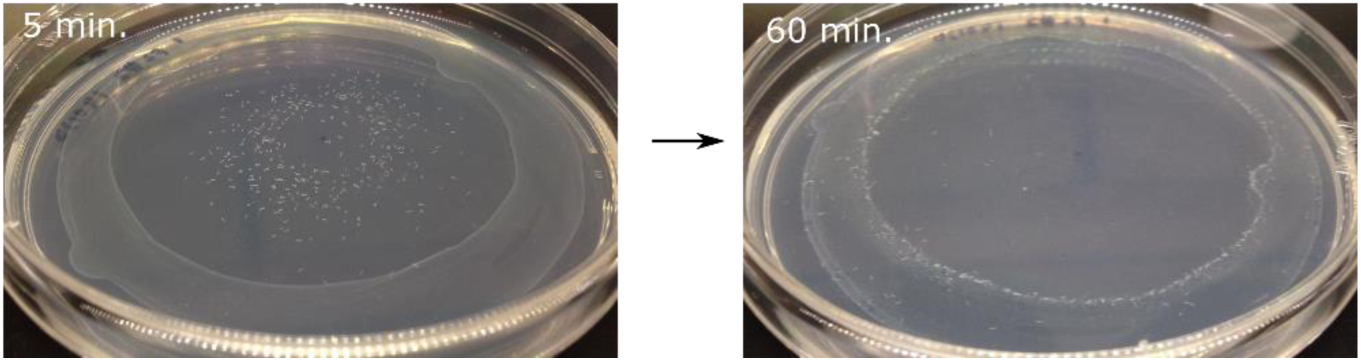
Photos of Edge Assay. (*Left*) Photo of Edge Assay after 5 min with worms moving away from the center. (*Right*) Photo of Edge Assay after 60 min with worms reaching and remaining in the edge.

**Table S1.** Number of screened genomes and isolated mutants.

|  | EMS-mutagenesis I |  |  | EMS-mutagenesis II |  |  | Total |
| --- | --- | --- | --- | --- | --- | --- | --- |
|  | Day 1 | Day 3 | Day 5 | Day 1 | Day 3 | Day 5 |  |
| **Genomes Screened** | 400 |  |  | 600 |  |  |  |
| **Isolated mutants** |  | 13 | 23 |  | 17 | 17 | 70 |
| **Viable isolated mutants** |  | 3 | 8 |  | 9 | 2 | 22 |
| **Isolated mutants with**<br>**reproducible deficits** |  | 0 | 3 |  | 2 | 0 | 5 |

**Figure S2.**
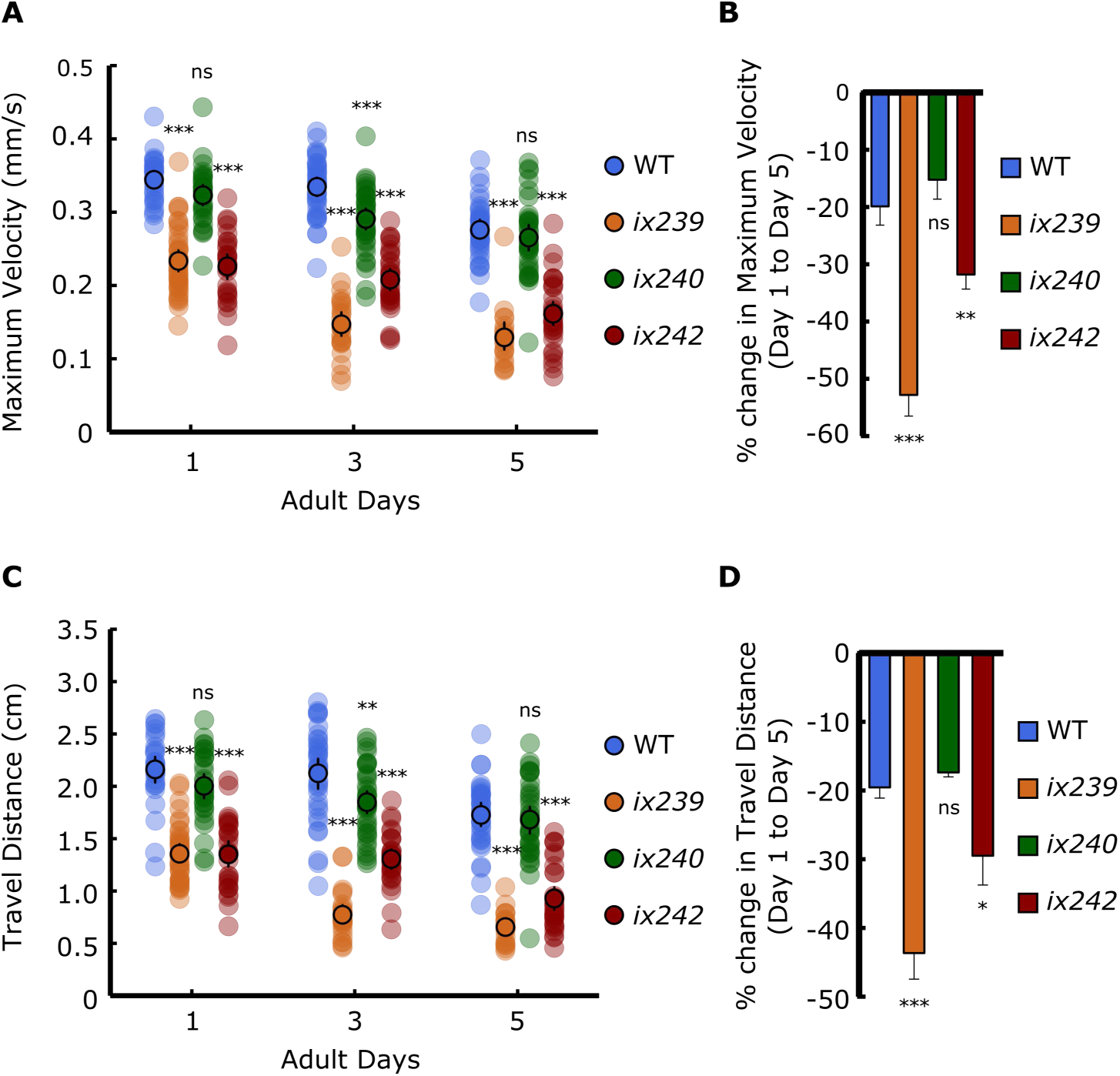
Maximum velocity and travel distance of isolated mutants. (A) Maximum velocities of WT, *ix239, ix240*, and *ix242* worms. (B) Percent change in maximum velocity of worms from A. (C) Travel distances of WT, *ix239, ix240*, and *ix242* worms. (D) Percent change in travel distance of worms from C. Error bars indicate 95% confidence intervals. For maximum velocity and travel distance experiments, n = 30–45 worms per strain for each day (10–15 worms from 3 biological replicate plates). For percent change in maximum velocity graphs, n = 3 biological replicate plates. \**P* < 0.05; \*\**P* < 0.01; \*\*\**P* < 0.001; ns, not significant; One-way ANOVA with Dunnett’s post hoc test vs. WT.

**Figure S3.**
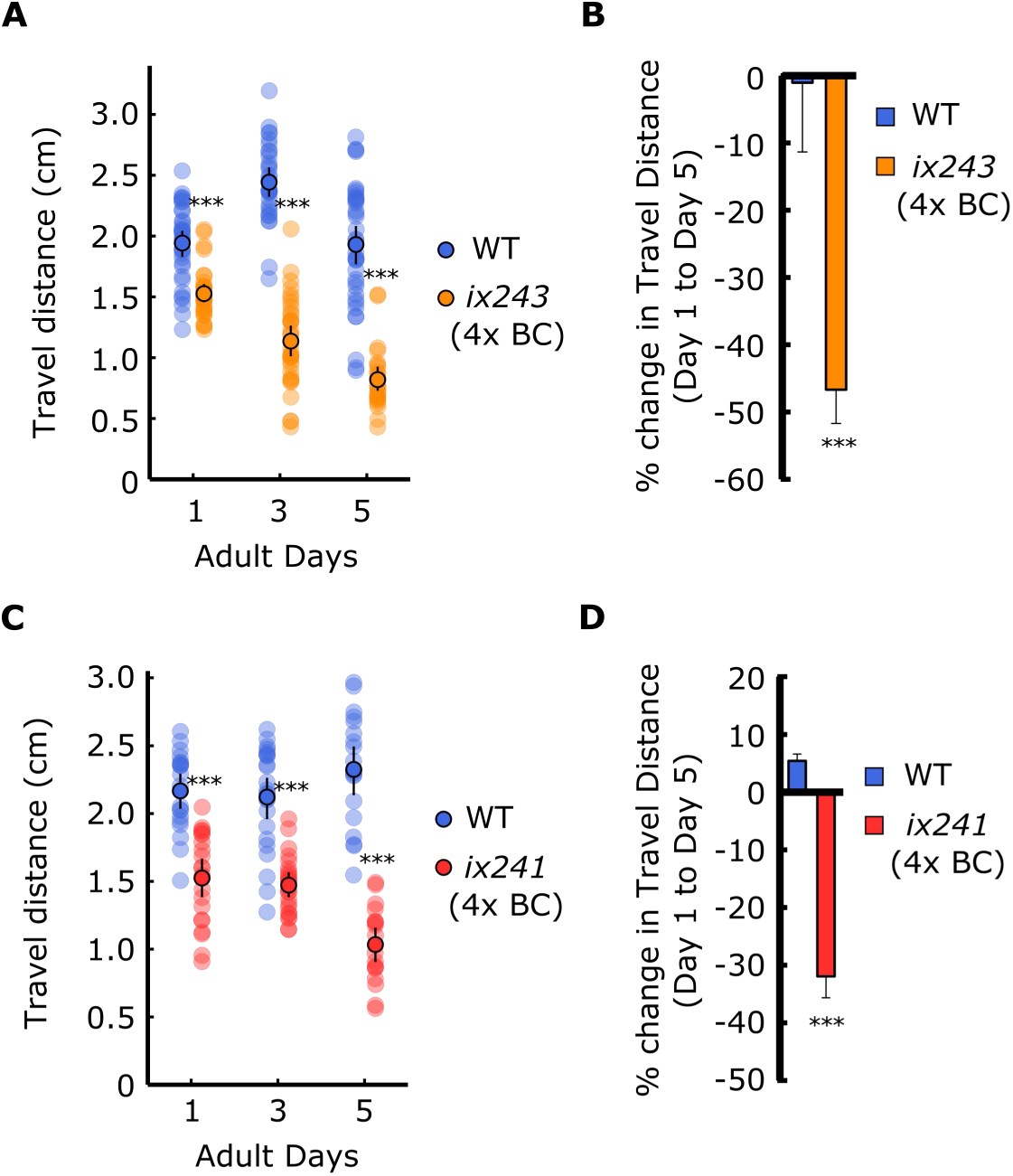
*ix241* and *ix243* worms show progressive locomotor decline after four backcrosses. (A) Travel distances of WT and *ix243* worms (backcrossed four times (4x BC)). (B) Percent change in travel distance of WT and *ix243*(4x BC) worms. (C) Travel distances of WT and *ix241*(4x BC) worms. (D) Percent change in travel distance of WT and *ix241*(4x BC) worms. For travel distance experiments, n = 30–45 worms per strain for each day (10–15 worms from 3 biological replicate plates). For percent change in travel distance graphs, n = 3 biological replicate plates. \*\*\**P* < 0.001; Unpaired Student’s *t* test vs. WT.

**Table S2.** Lifespan Analysis.

| **Strain** | **Median Lifespan (adult days)** | ***P* value vs WT (Log-rank test)** | **Worms (Counted/Total)** | **FUDR** |
| --- | --- | --- | --- | --- |
| N2 | 18 |  | 56/90 | 25 µg/ml |
| *ix243*(4x backcrossed) | 16 | *P* = 0.00078 | 89/90 | 25 µg/ml |
| N2 | 18 |  | 82/90 | 25 µg/ml |
| *ix243*(4x backcrossed) | 16 | *P* < 0.0001 | 84/90 | 25 µg/ml |
| N2 | 18 |  | 87/90 | 25 µg/ml |
| *ix243*(4x backcrossed) | 16 | *P* < 0.0001 | 87/90 | 25 µg/ml |
| N2 | 17 |  | 94/120 | 0 |
| *ix241*(4x backcrossed) | 16 | *P* = 0.095 | 77/120 | 0 |
| N2 | 14 |  | 66/90 | 0 |
| *ix241*(4x backcrossed) | 15 | *P* = 0.12 | 64/90 | 0 |
| N2 | 12 |  | 67/90 | 0 |
| *ix241*(4x backcrossed) | 14 | *P* = 0.024 | 74/90 | 0 |

**Figure S4.**
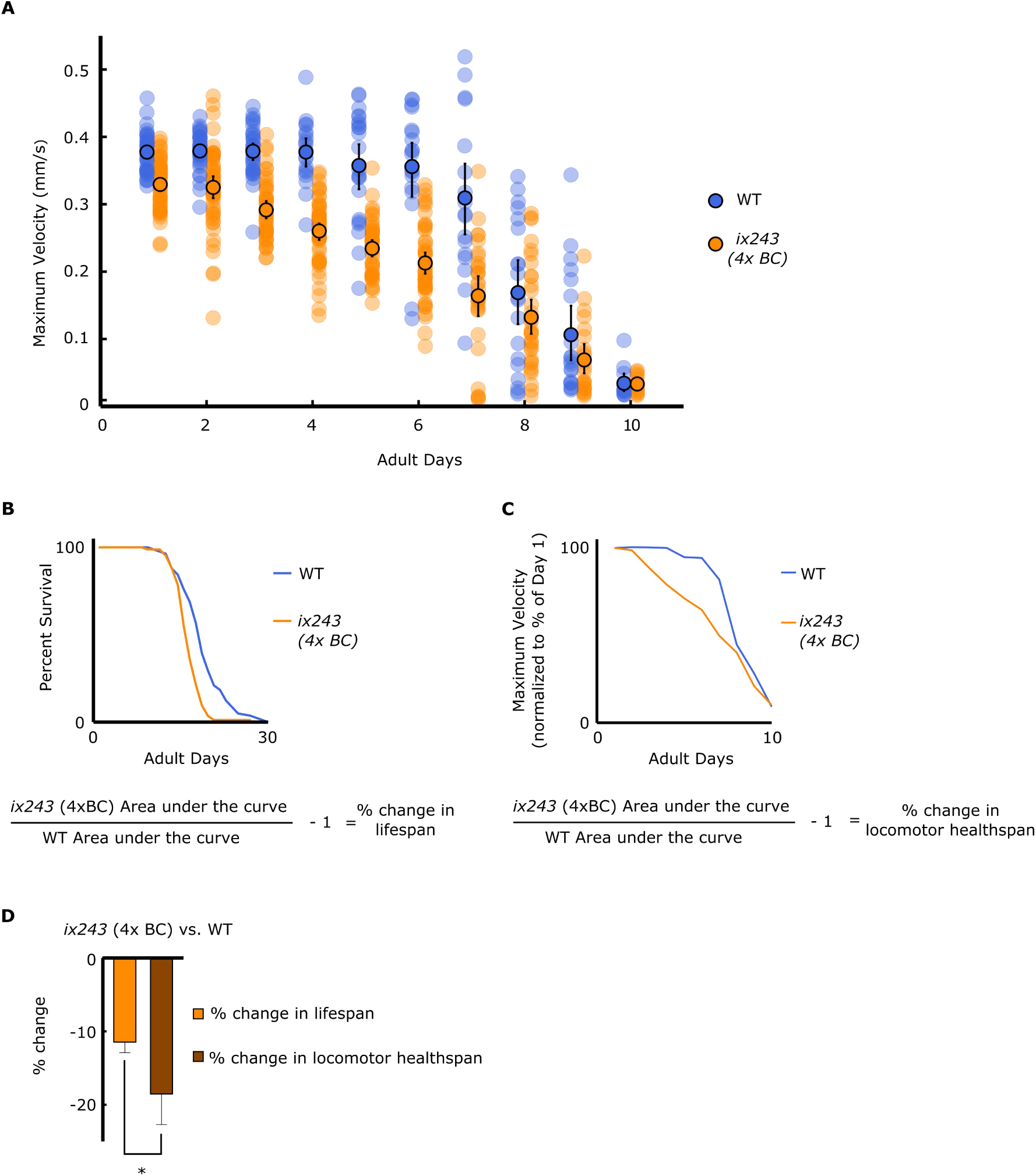
Greater reduction in total locomotor healthspan compared to lifespan in *ix243* worms. (A) Maximum velocities of WT and *ix243* (4x backcrossed (4x BC)) worms. n = 30–45 worms per strain for each day (10–15 worms from 3 biological replicate plates). (B) (Top) Representative survival curve of WT (n = 56 worms) and *ix243*(4x BC) worms (n = 89 worms). (Bottom) Calculation method of percent change in lifespan. (C) (Top) Representative decline in maximum velocity curve of WT and *ix243*(4x BC) worms. n = 30–45 worms per strain for each day (10–15 worms from 3 biological replicate plates). (Bottom) Calculation method of percent change in locomotor healthspan. (D) Percent change in lifespan (n = 3 biological replicate plates for WT and *ix243*(4xBC)) and locomotor healthspan (n = 3 biological replicate plates for WT and *ix243*(4xBC)) of *ix243*(4xBC) worms compared to WT. \**P* < 0.05; Unpaired Student’s *t* test.

**Table S3.** Development times of isolated mutant strains. Development time from egg to first egg-lay (n=5 worms per strain).

| Strain | Development Time (h) | % of WT |
| --- | --- | --- |
| WT | 70.4 | 100.0% |
| *ix239* | 74.4 | 105.7% |
| *ix240* | 71.2 | 101.1% |
| *ix241* (4x backcrossed) | 73.2 | 104.0% |
| *ix242* | 71.8 | 102.0% |
| *ix243* (4x backcrossed) | 80.2 | 113.9% |

**Figure S5.**
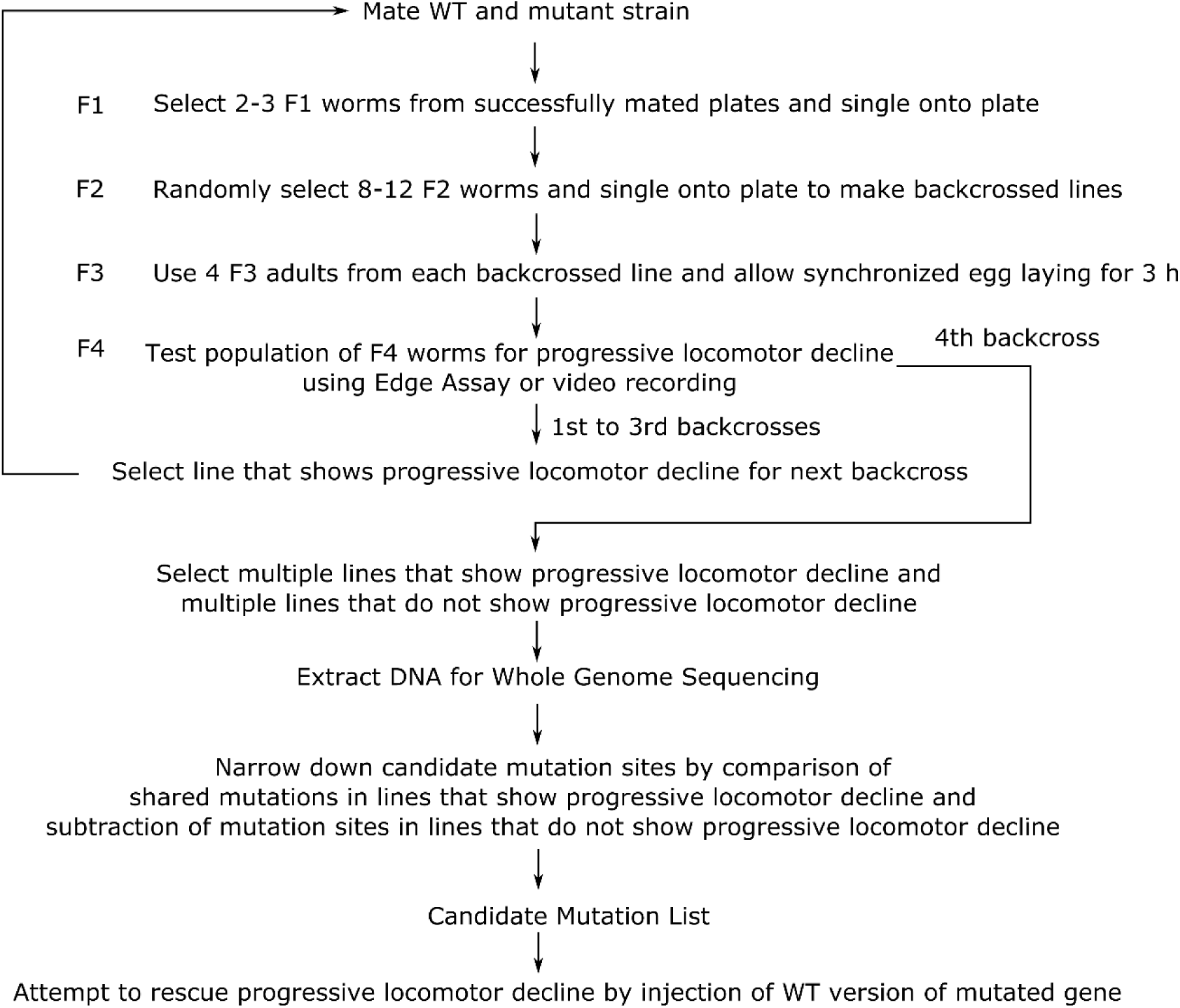
Strategy to identify causative mutation site.

**Table S4.**
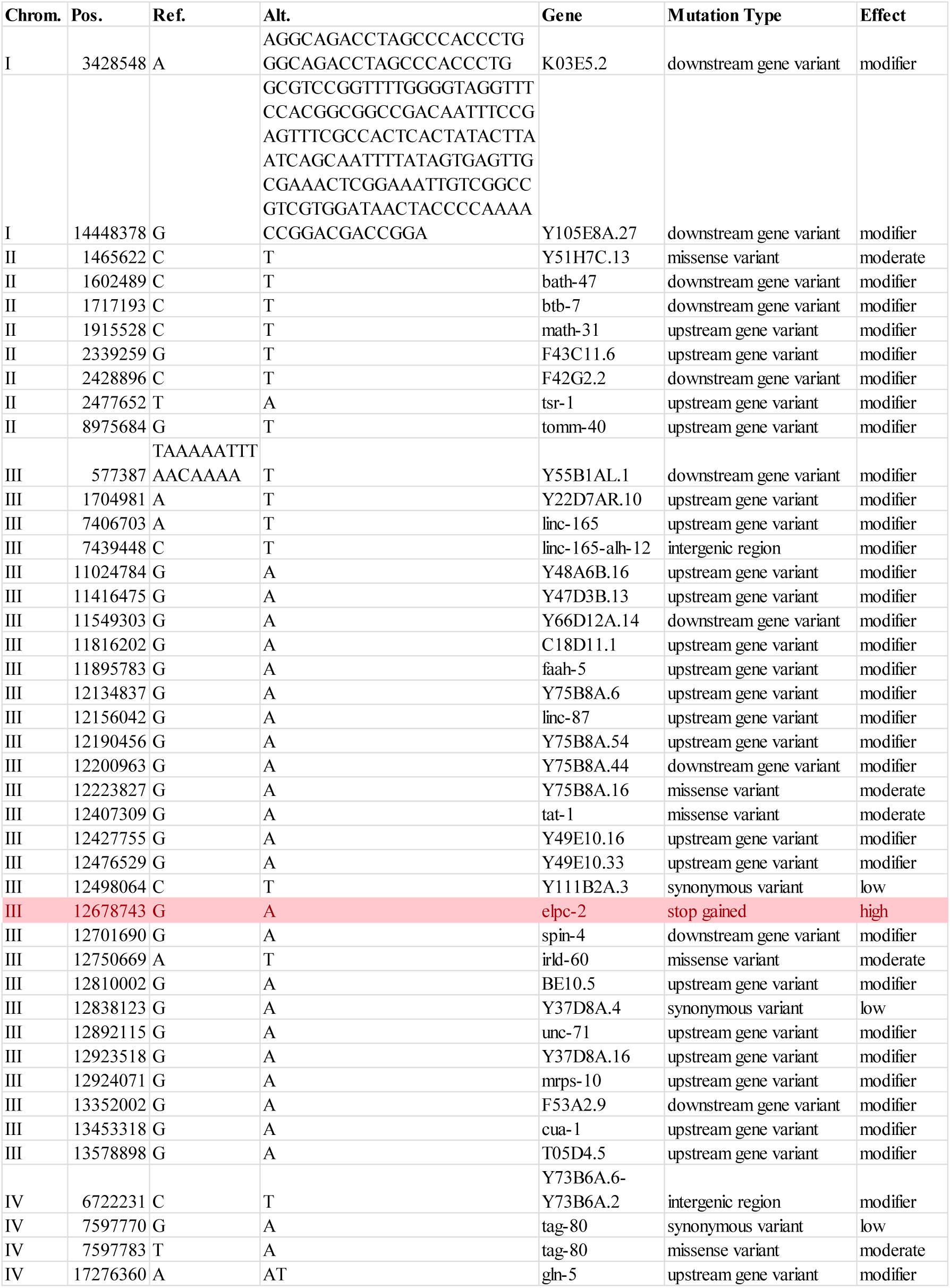
Remaining mutations in *ix243* mutant strains.

**Figure S6.**
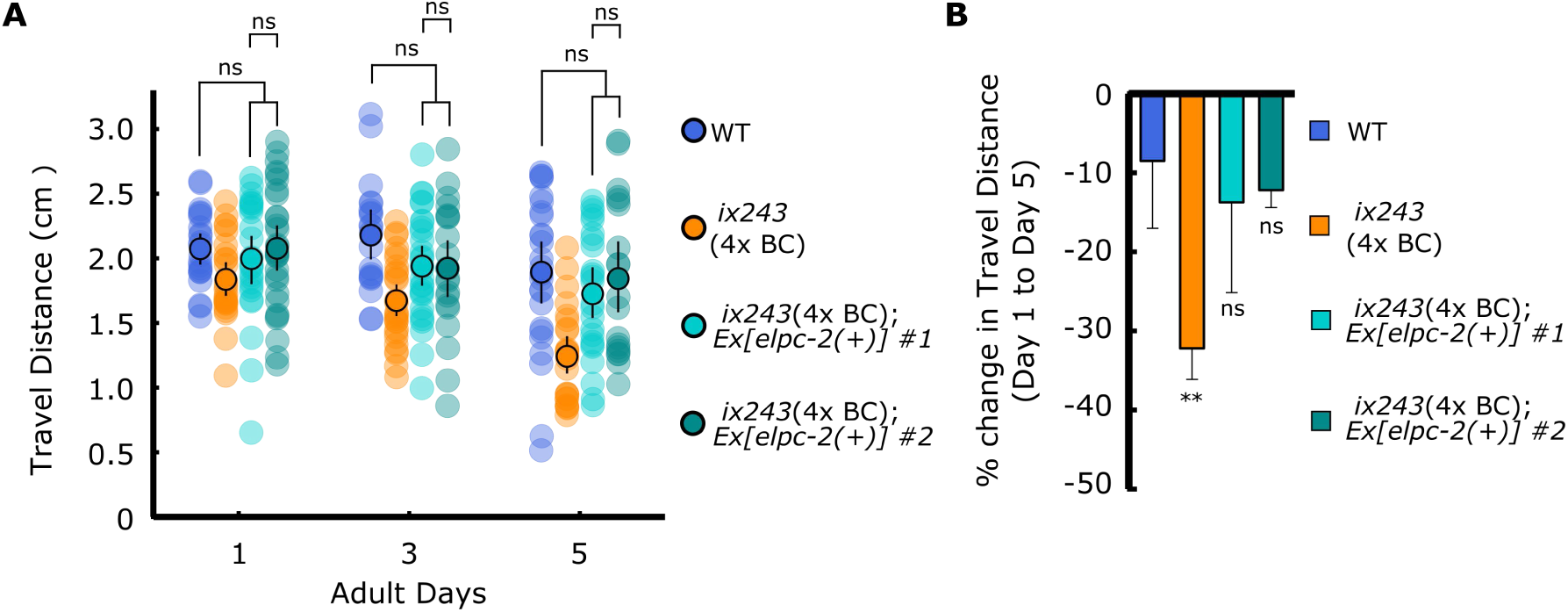
Wild-type *elpc-2* gene rescues progressive decline in locomotor function of the *ix243* mutant strain. (A) Travel distance of WT, *ix243*(4x BC), *ix243*(4x BC);*Ex[elpc-2(+)] #1, and ix243*(4x BC);*Ex[elpc-2(+)] #2* worms. n = 30–45 worms per strain for each day (10–15 worms from 3 biological replicate plates). (B) Percent change in travel distance of worms from A. n = 3 biological replicate plates. \*\**P* < 0.01; \*\*\**P* < 0.001; ns, not significant; One-way ANOVA with Dunnett’s post hoc test vs. WT.

**Figure S7.**
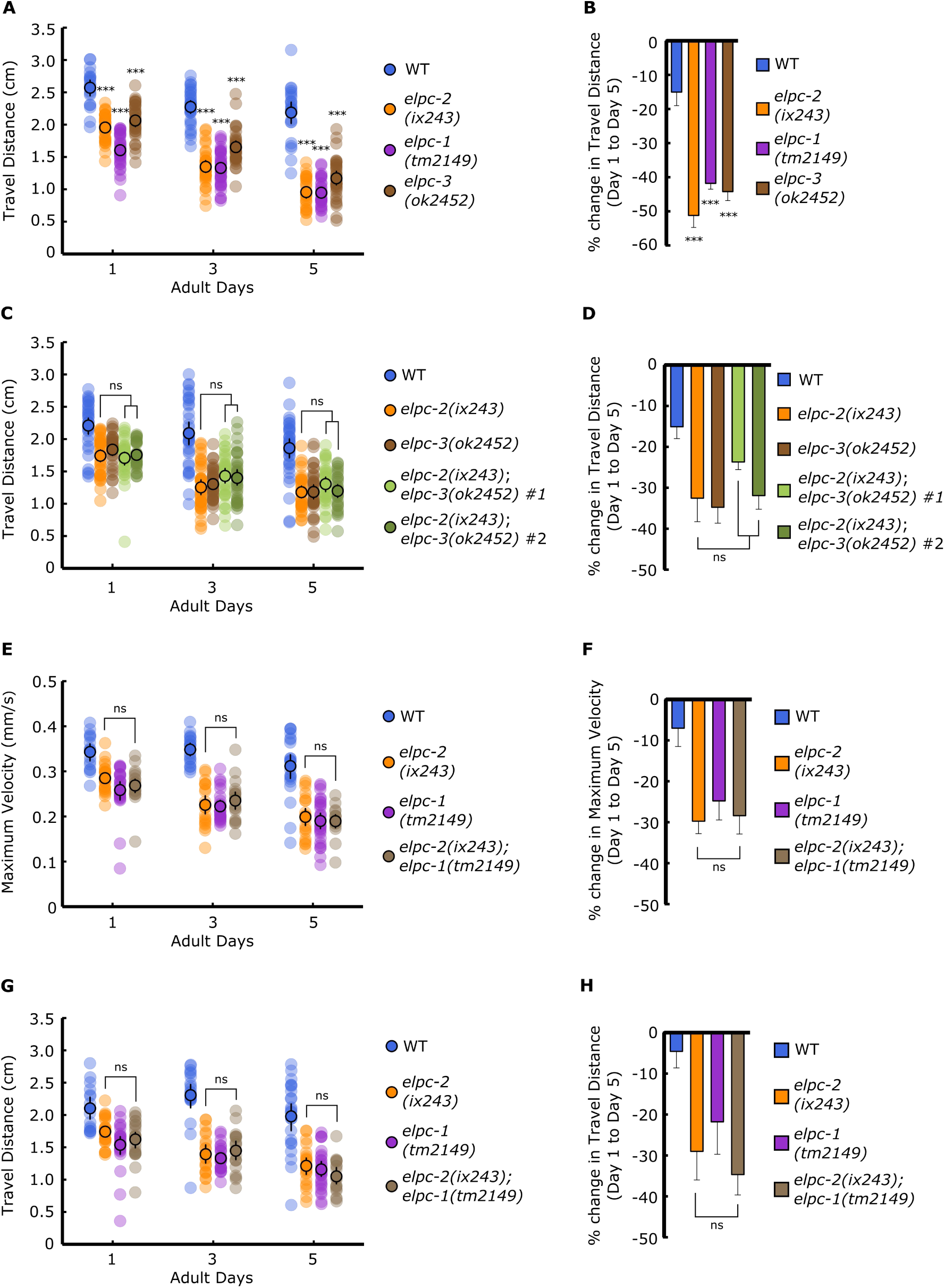
The Elongator complex is required to maintain adult locomotor function. (A) Travel distances of WT, *elpc-1(tm2149)* and *elpc-3(ok2452)* worms. (B) Percent change in travel distance of worms from A. (C) Travel distances of WT, *elpc-2(ix243)*, *elpc-3(ok2452)*, and *elpc-2(ix243);elpc-3(ok2452)* worms. (D) Percent change in travel distance of worms from C. (E) Maximum veolcities of WT, *elpc-2(ix243), elpc-1(tm2149), and elpc-1(tm2149);elpc-2(ix243)* worms. (F) Percent change in maximum velocity of worms from E. (G) Travel distances of WT, *elpc-2(ix243), elpc-1(tm2149), and elpc-1(tm2149);elpc-2(ix243)* worms. (H) Percent change in travel distance of strains from G. For maximum velocity and travel distance experiments, n = 30–45 worms per strain for each day (10–15 worms from 3 biological replicate plates). For percent change in maximum veloity graphs, n = 3 biological replicate plates. \*\*\**P* < 0.001; ns, not significant; One-way ANOVA with Dunnett’s post hoc test vs. WT for A, B; One-way ANOVA with Tukey’s post hoc test for C–H.

**Figure S8.**
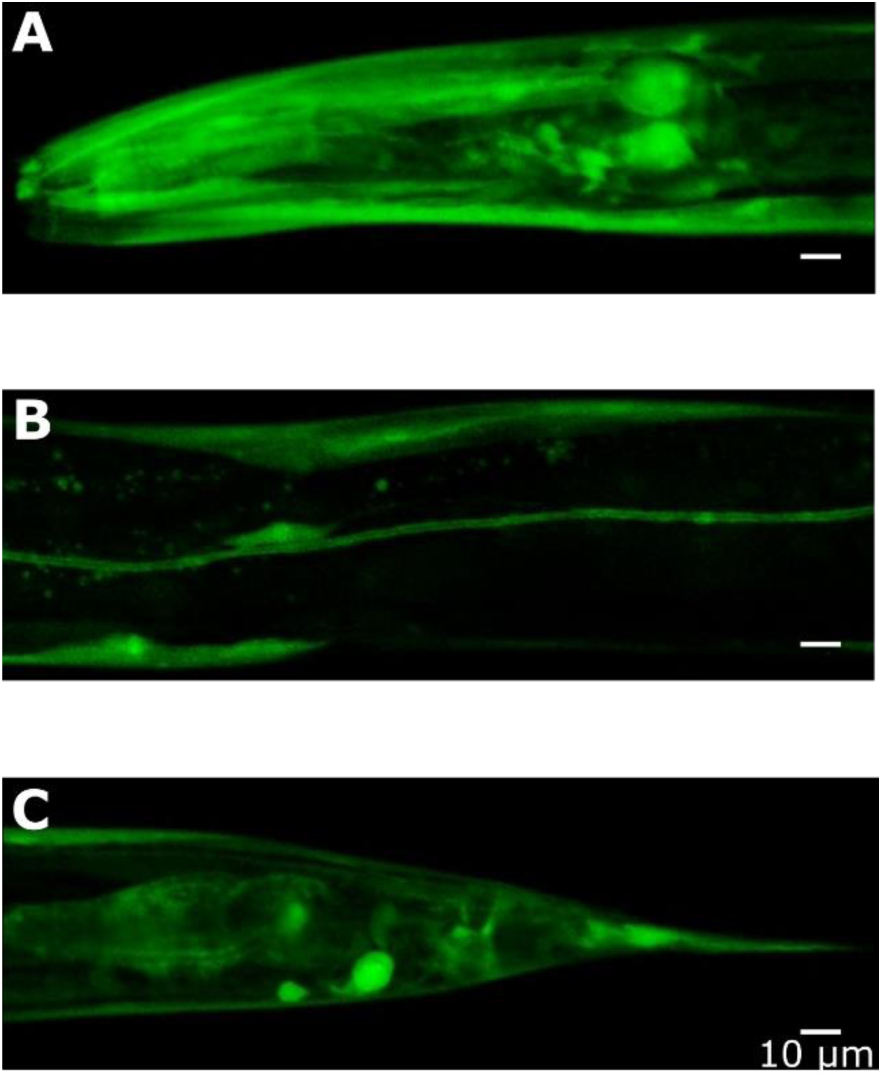
Expression pattern of *elpc-2* transcriptional GFP fusion. (A) *elpc-2p::GFP* expression in pharynx, neurons, and head muscles. (B) *elpc-2p::GFP* expression in body wall muscles and canal cell. (C) *elpc-2p::GFP* expression in coelomocytes, intestine, and tail. Scale bars: 10 µm.

**Figure S9.**
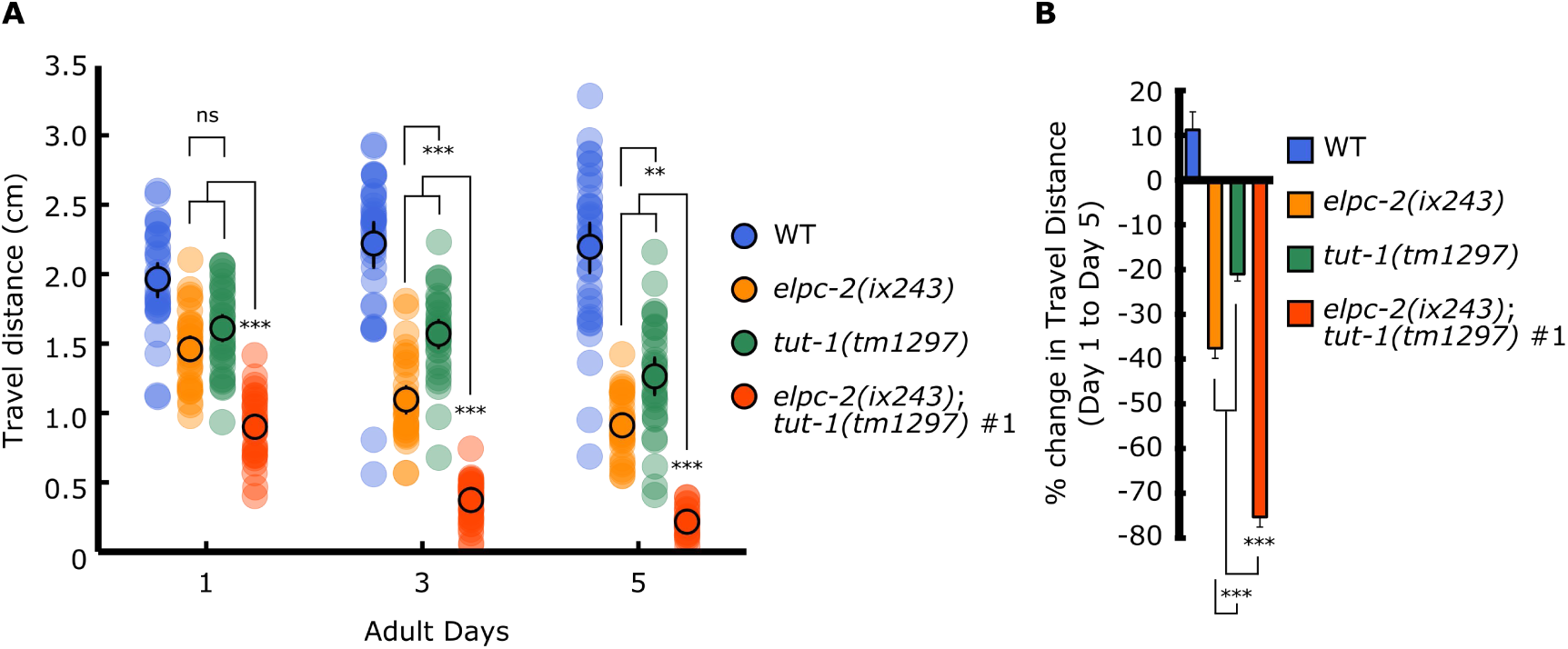
*tut-1(tm1297)* mutant shows progressive decline in locomotor function. (A) Travel distances of WT, *elpc-2(ix243)*, *tut-1(tm1297)*, and *elpc-2(ix243);tut-1(tm1297)* worms. n = 30–45 worms per strain for each day (10–15 worms from 3 biological replicate plates). (B) Percent change in travel distance of worms from A. Error bars indicate 95% confidence intervals. n = 3 biological replicate plates. \**P* < 0.05; \*\**P* < 0.01; ***P < 0.001; ns, not significant; One-way ANOVA with Tukey’s post hoc test.

**Table S5.** Development times of *tut-1(tm1297) and elpc-2(ix243);tut-1(tm1297)* mutants. Development time from egg to first egg-lay (n = 5 worms per strain).

| Strain | Development Time (h) | % of WT |
| --- | --- | --- |
| WT | 70.4 | 100.0% |
| *tut-1(tm1297)* | 82.0 | 116.5% |
| *elpc-2(ix243);tut-1(tm1297)* | 145.4 | 206.5% |

**Figure S10.**
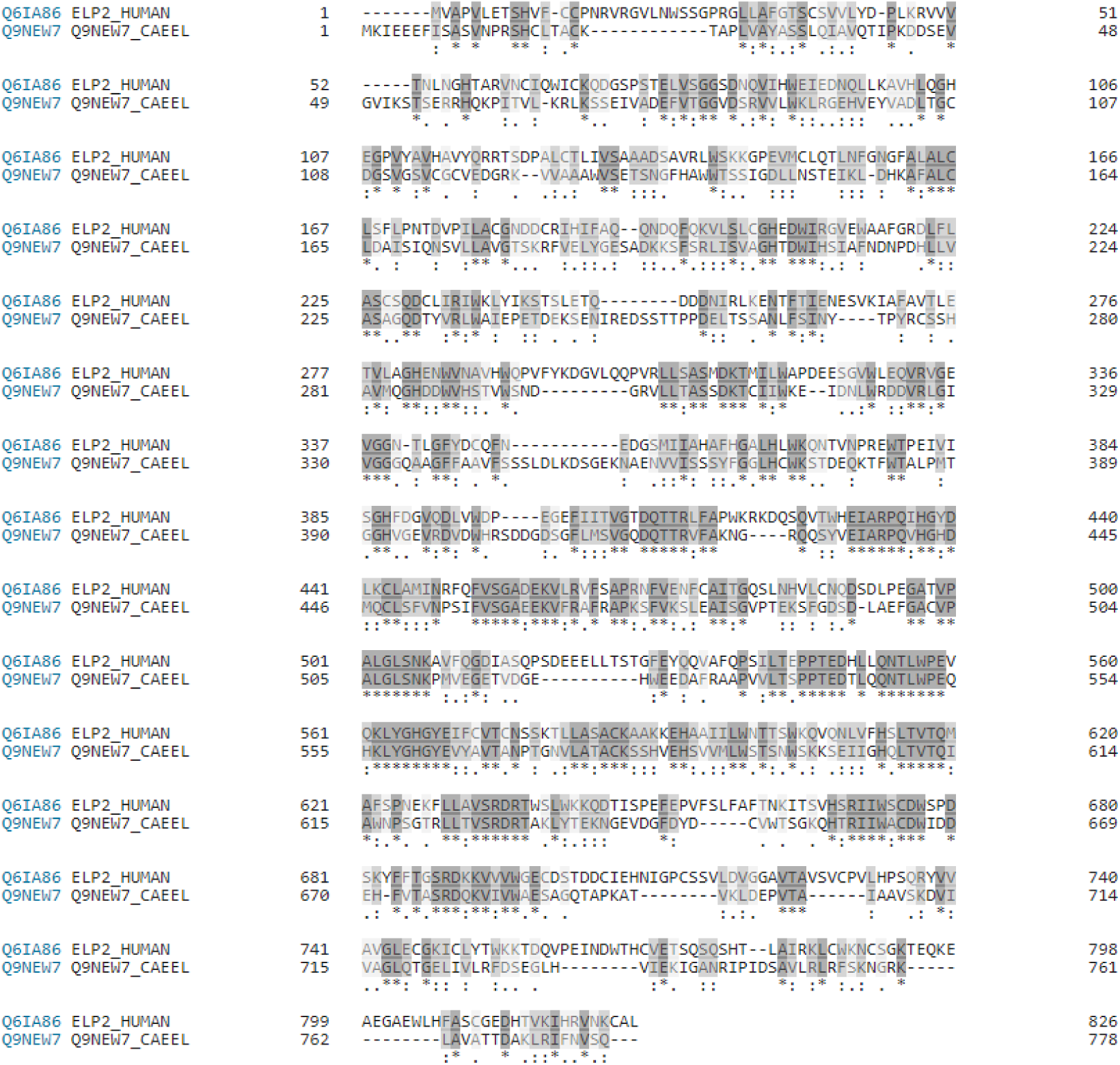
Amino acid alignment of *C. elegans* ELPC-2 and human ELP2. *C. elegans* ELPC-2 and human ELP2 were aligned using Clustal Omega (Sievers et al., 2011). An asterisk (*) indicates positions that are conserved; colon (:) indicates positions that are strongly similar (> 0.5 in the Gonnet PAM 250 matrix); period (.) indicates positions that are weakly similar (< 0.5 in the Gonnet PAM 250 matrix).

## Supplementary Information: Strain list

| **Strain** | **Genotype** | **Obtained From** |
| --- | --- | --- |
| CB408 | *unc-43(e408) IV* | CGC |
| CB190 | *unc-54(e190) I* | CGC |
| MT7929 | *unc-13(e51)* | CGC |
| AM725 | rmIs290 [*unc-54p* ::Hsa-sod-1 (127X)::YFP] | CGC |
| VC1937 | *elpc-3(ok2452) V* | CGC |
| CF1038 | *daf-16(mu86) I* | CGC |
| n/a | *elpc-1(tm2149) I* | National BioResource Project (Japan) |
| n/a | *tut-1(tm1297) IV* | National BioResource Project (Japan) |
| OF1260 | *ix239* | This study |
| OF1261 | *ix240* | This study |
| OF1262 | *ix241* | This study |
| OF1263 | *ix241* (4x backcrossed) | This study |
| OF1264 | *ix242* | This study |
| OF1265 | *elpc-2(ix243) III* | This study |
| OF1266 | *elpc-2(ix243) III* (4x backcrossed) | This study |
| OF1267 | *elpc-2(ix243) III;ixEx244[elpc-2(+);lin-44p::RFP]* | This study |
| OF1268 | *elpc-2(ix243) III;ixEx245[elpc-2(+);lin-44p::RFP]* | This study |
| OF1269 | *elpc-2(ix243) III;ixEx246[lin-44p::RFP]* | This study |
| OF1270 | *ixEx247[lin-44p::RFP]* | This study |
| OF1271 | *elpc-1(tm2149) I;elpc-2(ix243) III* | This study |
| OF1272 | *elpc-2(ix243) III;elpc-3(ok2454) V (#1)* | This study |
| OF1273 | *elpc-2(ix243) III;elpc-3(ok2454) V (#2)* | This study |
| OF1289 | *elpc-2(ix243) III;tut-1(tm1297) IV* | This study |

## Supplementary Information: Primer list

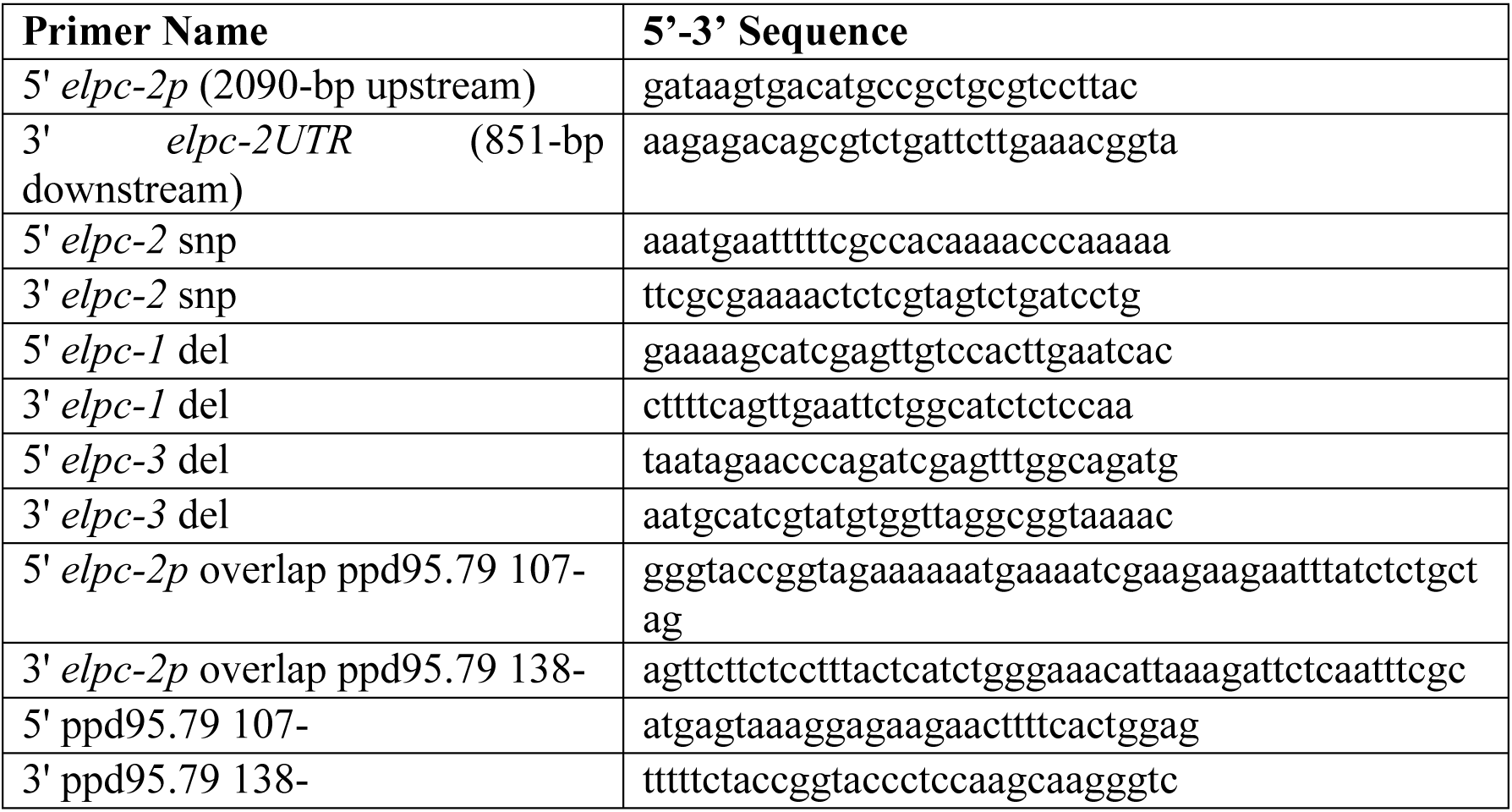

